# Striatal Social Reward Sensitivity Predicts Trust-Related Brain Responses Depending on Closeness and Depression

**DOI:** 10.64898/2026.03.27.714332

**Authors:** Shenghan Wang, Yi Yang, Cooper J. Sharp, Dominic Fareri, Jason Chein, David V. Smith

## Abstract

**Background:** Depression is associated with social dysfunction, but the mechanisms linking affective symptoms to disrupted close relationships remain poorly understood. One possibility is that depression alters how people experience rewards shared with close others and how they interpret partners’ actions. It remains unclear whether neural sensitivity to shared reward predicts social valuation during more complex interactions such as reciprocated trust.

**Methods:** In this preregistered fMRI study, participants completed a reward-sharing task and a Trust Game with a close friend, a stranger, and a computer. We measured striatal shared reward sensitivity (SRS; friend > computer) and tested whether it related to subsequent investment behavior and brain responses to trust reciprocation. Depressive symptoms and perceived closeness were assessed via self-report.

**Results:** In a final sample of n = 123, participants reporting more depressive symptoms invested more in their friend than in the computer. Striatal SRS predicted temporoparietal junction responses to reciprocated trust, but this association depended jointly on social closeness and depression — with depression reversing the expected pattern among individuals reporting closer relationships. Striatal SRS was also inversely associated with connectivity between the default mode network and cerebellum during reciprocity.

**Conclusions:** These findings suggest that closeness calibrates the striatal SRS’ link to regional activity and network-level responses during social exchange, while depression alters how striatal SRS relates to regional activity, potentially disrupting how individuals interpret and respond to close others.

## 1. Introduction

Real-life decisions are often made in socially salient contexts, where choices are shaped by the actions of and effects on others. A question of broad interest is how the social environment affects reward valuation signals that guide choice behavior (Lee & Chung, 2022), and whether perceived social connectedness with those involved modulates this influence — potentially leading to adaptive or maladaptive decision making. These interactions may be further shaped by internal states that influence how individuals represent both reward value and social meaning. Depression, in particular, is known to disrupt both reward circuitry (Ng et al., 2019) and social information processing (Hames et al., 2013; Weightman et al., 2014), with effects that can persist even after clinical remission (Kennedy et al., 2007; Rhebergen et al., 2010). Further, reward and social dysfunction may intermingle in close relationships, where diminished feelings of social closeness have been reported in depression (Frick et al., 2021).

Although reward valuation, social context processing, and representations of social closeness are related, converging evidence suggests they rely on partially distinct neural systems: social reward valuation is closely tied to ventral striatal circuitry, whereas representations of social closeness are more consistently associated with the default mode network (DMN), including mPFC and TPJ. Consistent with this distinction, the ventral striatum (VS) preferentially activates during rewarding experiences involving close others, such as adolescent decision making in the presence of peers (Chein et al., 2011), sharing monetary rewards with a close friend (Fareri et al., 2012), and when a trusted friend reciprocates monetary investment (Fareri et al., 2015). In contrast, social closeness is more typically reflected in activity within core DMN regions in social perception (e.g., Roseman-Shalem et al., 2024) and social valuation tasks (e.g., Fareri et al., 2020), particularly where the DMN overlaps with the "social brain" (Shultz & Dunbar, 2012), including medial prefrontal cortex (mPFC, Krienen et al., 2010) and temporoparietal junction (TPJ, Carter & Huettel, 2013). It remains unknown, however, whether VS responses to social valuation generalize across contexts and predict the neural substrates of social closeness perception in other tasks.

Despite these dissociations, disruptions in both reward processing and social functioning in depression suggest that the link between social closeness representations and reward valuation may be altered. Depression is thought to be rooted in aberrant reward valuation and motivational processes though recent work has revealed additional nuance (Chung et al., 2017; Heffner et al., 2021), with blunted striatal responding long documented as a key neural underpinning of anhedonia (Cooper et al., 2018; Forbes et al., 2009; Pizzagalli et al., 2008). Social-domain specific anhedonia and social withdrawal are frequently reported symptoms as well (Blanchard et al., 2001; Kuan et al., 2024; Setterfield et al., 2016), and aberrant DMN activation has been observed in depressive patients (Sheline et al., 2009), while a reduced DMN intrinsic connectivity has been observed in those with greater social dysfunction (Saris et al., 2020). Consistent with this observation, studies of maternal depression report altered motivation and neural responses to infant cues, including reduced striatal and mPFC activation (Laurent & Ablow, 2012, 2013; Macrae et al., 2015).

How social context processing is altered in depression remains inconclusive, with comparable numbers of studies reporting blunting (Ait Oumeziane et al., 2019; Caouette & Guyer, 2016; Gradin et al., 2015) or sensitization (Gillard et al., 2021; Healey et al., 2014; Sankar et al., 2019; Steger & Kashdan, 2009; Yttredahl et al., 2018) to social outcomes in depression. This inconsistency may reflect varied experimental manipulations of social context and uncontrolled salience of social relationships in studies where perceived closeness has not been explicitly measured. Studies also vary in whether they assess passive responses to social feedback versus valuation processes that guide behavior under uncertainty. Behavioral economic games offer a useful way to probe these issues, linking social valuation to observable decisions while allowing rigorous experimental control over social outcomes (Vanyukov et al., 2019).

This study examines whether VS responses to social context and closeness during a reward task predict cross-task behavior and brain function, and whether depression alters that link. Participants completed a card-guessing task involving reward sharing and a Trust Game involving investment under the risk of defection, with the same partners across both tasks. Prior work has operationalized social closeness in different ways—comparing social partners at varying levels of perceived closeness (e.g., Roseman-Shalem et al., 2024) or social partners with nonsocial agents or objects (e.g., Baetens et al., 2014). Notably, these contrasts isolate overlapping but distinct aspects of social reward processing: a friend–computer contrast captures the combined influence of social presence and relational closeness, whereas a friend–stranger contrast holds social presence constant to isolate closeness-specific variation (Fareri et al., 2012; Smith et al., 2026). Thus, we preregistered analyses focusing on the friend–computer contrast as a test of socially shared reward relative to a nonsocial control, with friend–stranger contrasts as a secondary test of how relational closeness modulates reward sensitivity. We tested activation and connectivity patterns involving VS, mPFC, TPJ, and DMN, asking three key questions. First, does striatal shared reward sensitivity (SRS) predict behavior and brain activation in the Trust Game? Second, are these effects moderated by perceived closeness and depressive symptoms? Finally, do these relationships extend to functional connectivity between the DMN and key social valuation regions?

## 2. Methods and Materials

### 2.1 Participants

We recruited 225 participants from the greater Philadelphia area (preregistration: https://osf.io/p654khttps://osf.io/p654k￼). All provided written informed consent under Temple University IRB-approved protocol. Data were collected as part of a larger project (Smith et al., 2024) and are publicly available on OpenNeuro￼https://openneuro.org/datasets/ds005123/versions/1.1.3￼). After preregistered exclusions (detailed in Supplementary Methods), final samples of n = 123 were retained for all analyses (demographic details in Supplementary Table 1). A G*Power sensitivity analysis (α=.05, 1−β=.80) indicated that with n=123 our preregistered interaction test was calibrated to detect effects of f²≥0.09 with 80% power (smaller effects would have lower power).

### 2.2 Experimental tasks

Participants performed two tasks in PsychoPy (Peirce et al., 2019) in counterbalanced order: a Shared Reward Task and a Trust Game, each involving social and nonsocial partners (friend, stranger, or computer). Participants provided a real-life friend’s photo and were told their friend would serve as one of their partners. To ensure incentive compatibility, participants were informed that bonus payments were determined by outcomes from one randomly selected trial per task.

In the Shared Reward Task (Figure 1a), participants guessed whether a hidden number was greater or less than five. Correct guesses yielded a $10 gain for both the participant and the partner; incorrect guesses yielded a $5 loss for both. In the Trust Game (Figure 1b), participants received an $8 endowment and chose how much to invest with their partner; invested amounts were tripled, and partners either reciprocated (returning half) or defected (keeping all). All partners had a 50% reciprocation rate. For full task details, see the Supplementary Methods and the accompanying data descriptor Smith et al. (2024).

**Figure 1.**
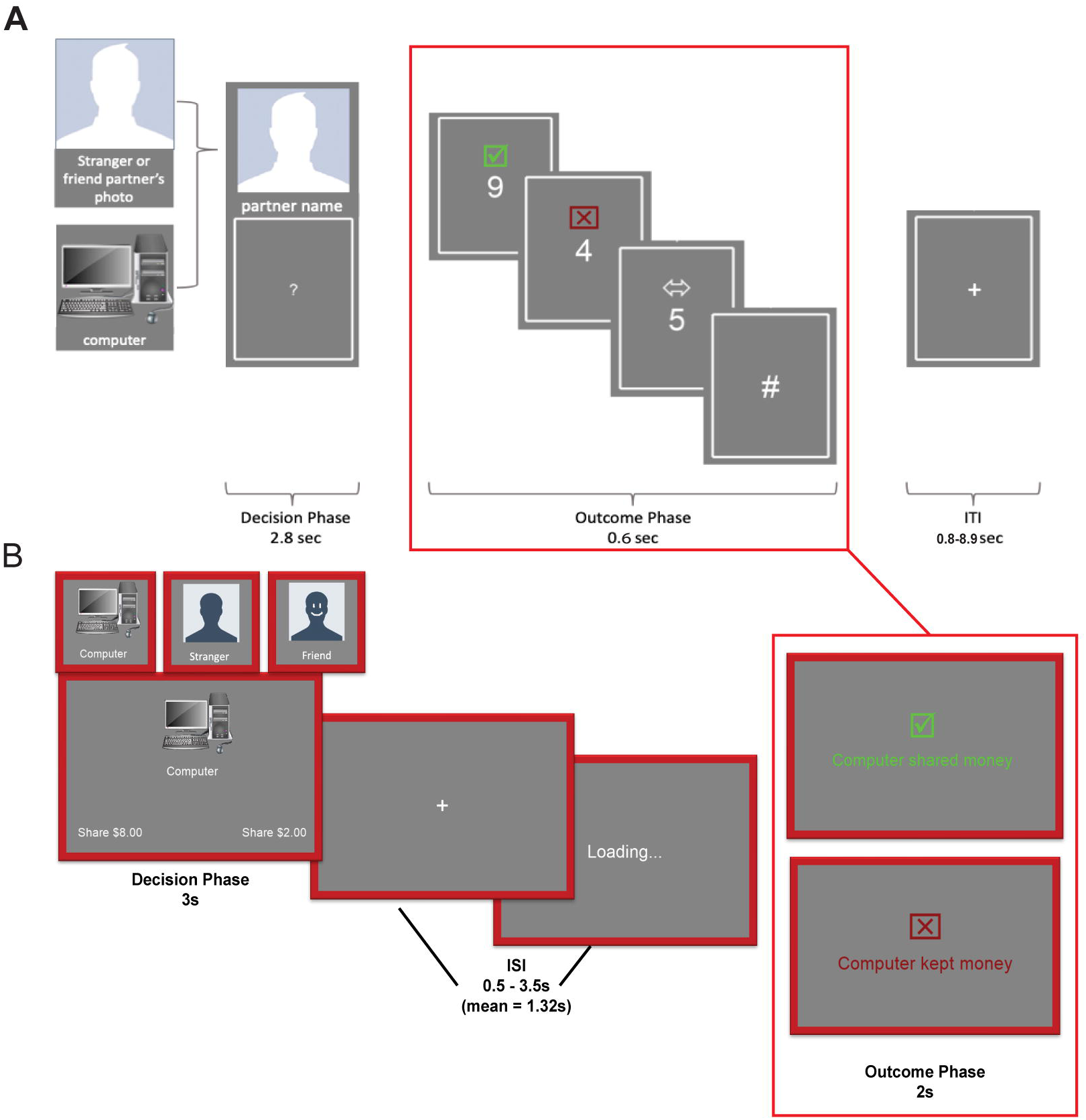
Study overview and task design. A) Shared Reward Task: On each trial, participants viewed one of three partners (friend, stranger, or computer) and guessed whether a randomly generated number (1–9) would be above or below 5. Correct guesses resulted in both the participant and partner winning $10; incorrect guesses resulted in a $5 loss for both; trials landing on 5 yielded no change. Striatal responses during reward > loss outcomes with a friend were extracted as the index of striatal shared reward sensitivity (SRS). B) Trust Game: Participants played with the same three partners. On each trial, they decided how much of an $8 endowment to invest. Investments were tripled, and the partner either reciprocated by returning half or defected by keeping the full amount. striatal SRS was used to predict neural responses during the outcome phase of trials in which a friend reciprocated trust. Figures adapted from Smith et al. (2024) with permission.

### 2.3 Individual difference measures

Depression symptoms were assessed at the first appointment using the “Down” subscale of the 7 Up 7 Down Inventory (Youngstrom et al., 2013), a seven-item, four-point questionnaire measuring the frequency of depressive behaviors and feelings (e.g., “There have been times when you really got down on yourself and felt worthless”). Perceived social closeness toward each partner was assessed using the Inclusion of Other in Self (IOS) scale (Aron et al., 1992). Participants viewed a seven-point series of overlapping circles depicting increasing self–other overlap and selected the image that best represented their relationship with each partner (friend, stranger, and computer).

### 2.4 Imaging data acquisition and preprocessing

Imaging data were collected at the Temple University Brain Imaging and Research Center using a 3T Siemens PRISMA scanner and a 20-channel head coil. Functional images were collected using a multi-echo echo-planar imaging (EPI) sequence with four echoes (TEs = 13.8, 31.54, 49.28, 67.02 ms), a TR of 1615 ms, voxel size of 2.7 mm³, multiband acceleration (factor = 3), and GRAPPA = 2. A total of 51 axial slices were acquired with a 10% inter-slice gap. To support motion correction and registration, single-band reference images were collected for each run. For co-registration of the functional images, high-resolution T1-weighted anatomical images were acquired using an MPRAGE sequence (TR = 2400 ms, TE = 2.17 ms, flip angle = 8°, voxel size = 1 mm³). Additional MRI acquisition and preprocessing procedures are detailed in the supplement and the technical details of acquisition, including variations in flip angle and the inclusion of phase images, are described in Smith et al. (2024).

Before further processing, structural and functional data were converted to BIDS format using HeuDiConv (Halchenko et al., 2024). All data were then preprocessed using the default settings in fMRIPrep (version 24.1.1; Esteban et al., 2019), including skull stripping, spatial normalization to MNI space, slice-timing correction, and motion correction (details in Supplemental Methods). Field maps were created from the available phase images using the MEDIC algorithm (Van et al., 2023) and used for susceptibility distortion correction (Supplemental Methods). Following fMRIPrep, functional data were smoothed using a 5 mm full-width at half-maximum (FWHM) Gaussian kernel in FSL, and grand-mean intensity normalization was applied using a single multiplicative factor.

### 2.5 Neuroimaging Analyses

Neuroimaging analyses including activation and connectivity were performed on FSL version 6.0.7 (Jenkinson et al., 2012; S. M. Smith et al., 2004) using general linear models with local autocorrelation (Woolrich et al., 2001). First-level models for each task included outcome and decision phase regressors differentiated by partner identity, convolved with a double-gamma hemodynamic response function. To assess task-dependent functional connectivity, we used a network-based psychophysiological interaction (nPPI) approach with the default mode network (Smith et al., 2009) as seed, examining coupling with mPFC and VS during Trust Game outcomes. Full model specifications are provided in the Supplementary Methods.

After combining runs with a fixed-effects model, group-level analyses used FSL FLAME Stage 1 (Woolrich et al., 2004). Analyses followed a preregistered ROI-first approach before proceeding to whole-brain analyses. ROIs were defined a priori: mPFC from a Neurosynth association map (keyword "mPFC," z > 8), TPJ from the Mars connectivity atlas, and ventral striatum from the Oxford–GSK–Imanova Striatal Connectivity Atlas. We defined striatal SRS as VS activation in the contrast of reward > loss sharing outcomes with a friend vs. other partners. Whole-brain z-statistic maps were thresholded at z > 3.1 with cluster-extent correction at p < 0.05 (Worsley, 2001). Brain imaging results are displayed using MRIcroGL (Rorden, 2025).

### 2.6 Deviations from Pre-registration

Our pre-registered analyses focused on the contrast between close friends and computer partners (Friend > Computer) for testing hypotheses about activation in mPFC and TPJ and connectivity between the ventral striatum (VS) or mPFC and the default mode network (DMN) during trust reciprocation. One section of the preregistration refers to the contrast as Friend > Stranger, though all other sections clearly specify Friend > Computer. For consistency with our stated hypotheses and analytic plan, we interpret this as an isolated inconsistency and focus on the Friend > Computer contrast in the main text, with Friend > Stranger results reported in the Supplement. After the primary model investigating closeness’s effect on striatal SRS returned null effects, we conducted follow-up analyses not specified in the preregistration, specifically through reduced models without depression terms.

## 3. Results

### 3.1 Does Striatal SRS Predict Trust Behavior?

We first confirmed that participants rated their friend as significantly closer than the computer partner (β = –3.65, SE = 0.13, t(244) = –28.13, p < .0001; Figure 2a), and invested more in their friend than in the computer (β = 1.206, SE = 0.09, t(244) = 12.96, p < .0001; Figure 2b). We then tested our preregistered hypothesis that greater closeness to the friend would amplify VS shared-reward sensitivity — the rationale being that a stronger social bond might enhance the striatal SRS. In a model including closeness, depressive symptoms, and their interaction, VS shared-reward sensitivity was not significantly associated with closeness (β = –0.012, SE = 0.053, t = –0.23, p = .82), and this association was not moderated by depression (β = –0.003, SE = 0.008, t = –0.39, p = .70). Results were similar when using an alternative friend–stranger contrast (see Supplement). These findings suggest that VS shared-reward sensitivity operates independently of perceived relationship strength, reflecting a more general signal for social reward value.

**Figure 2.**
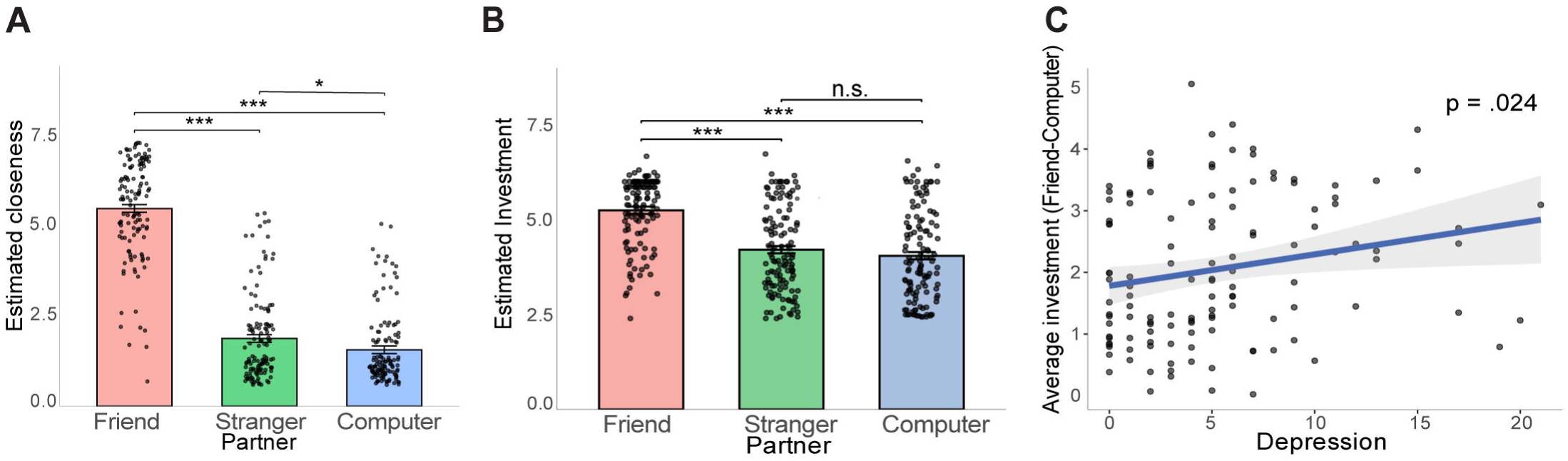
Partner closeness and depressive symptoms shape trust behavior. A) Participants rated their friend as significantly closer than the stranger or computer on the Inclusion of Other in Self scale. B) Participants invested more in their friend than in other partners. C) Higher depressive symptoms were associated with greater investment in the friend relative to the computer. Panels A and B show fixed-effect estimates from mixed-effects models with random intercepts, overlaid with raw data points. Panel C shows the association between depressive symptoms and partner-selective investment, controlling for age and gender. ***p < .001, *p < .05; error bars and shaded areas reflect standard error.

We next examined whether VS shared-reward sensitivity predicts trust behavior, operationalized as relative investment in the friend versus computer. VS shared-reward sensitivity was not associated with investment differences (β = –0.110, SE = 0.180, t = –0.61, p = .54), and this relationship was not moderated by closeness (β = 0.110, SE = 0.118, t = 0.93, p = .35) or predicted by closeness alone (β = 0.074, SE = 0.072, t = 1.02, p = .31). In contrast, individuals reporting more depressive symptoms invested more in their friend than in the computer (β = 0.051, SE = 0.022, t = 2.29, p = .024; Figure 2c). Together, these results indicate that VS shared-reward sensitivity did not account for individual differences in trust behavior, whereas depressive symptoms were associated with greater trusting investment to close partners.

### 3.2 Does Striatal SRS Predict Activation During Reciprocity?

To establish that our paradigm captured the expected social modulation of striatal reward responses, we first examined whether VS activation varied as a function of partner closeness across both tasks. As predicted, VS activation was strongest when outcomes were shared with or reciprocated by a close friend (Figure 3; see Supplementary Results S1 for full statistics), replicating prior work (Fareri et al., 2012; 2015). We then tested our preregistered hypothesis that individuals with greater VS shared-reward sensitivity would show increased activation in cortical regions implicated in social valuation and inference — specifically mPFC and TPJ — when a close friend reciprocated trust. Using preregistered ROI models, we tested whether striatal SRS predicted activation during reciprocate > defect outcomes with the friend relative to the computer. Across both regions, there were no significant associations with striatal SRS (mPFC: β = −0.404, SE = 0.302, t = −1.34, p = .18; TPJ: β = 0.035, SE = 0.240, t = 0.15, p = .89), and no evidence of moderation by closeness or depression. In a follow-up model including all interaction terms, a three-way interaction emerged in right TPJ (β = −0.027, SE = 0.013, t = −2.14, p = .035): greater striatal SRS was associated with increased TPJ activation during a friend’s reciprocation, but only among participants reporting high depressive symptoms and low closeness.

**Figure 3.**
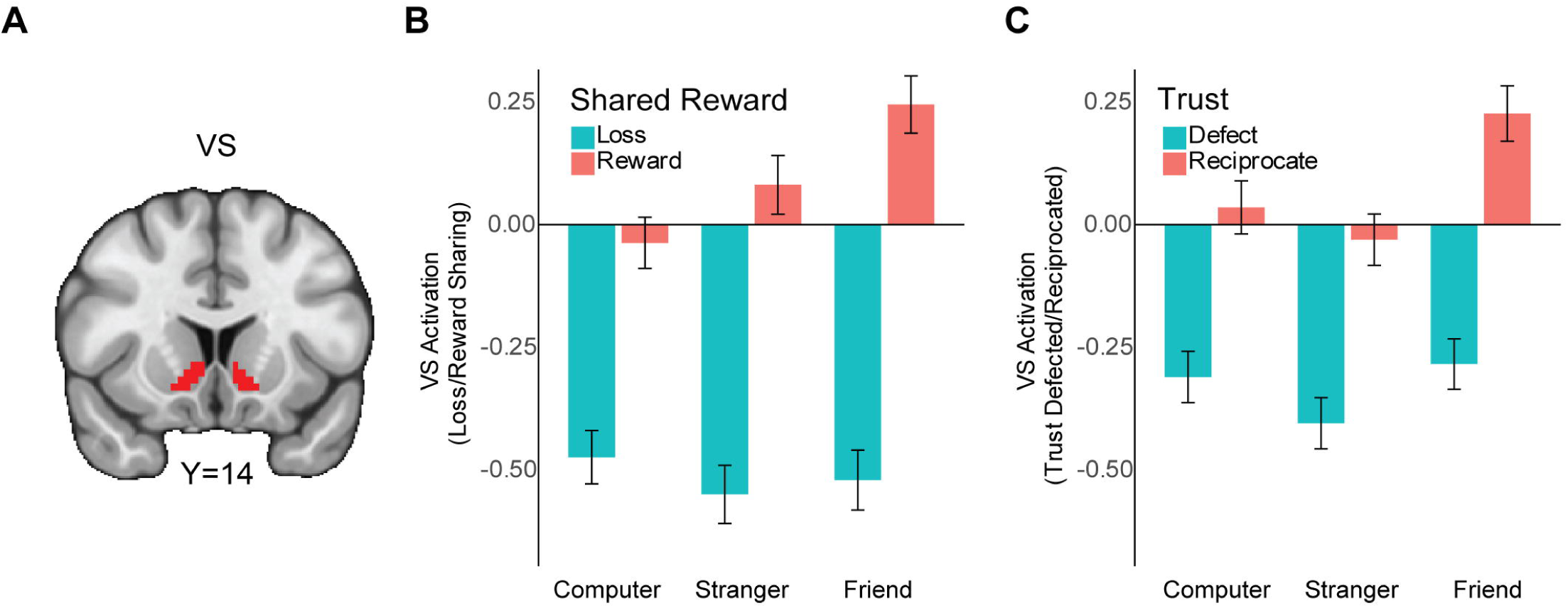
Ventral striatal activation is strongest during reward sharing and reciprocation with a friend, but does not differentiate partners during loss or defection. A) We conducted two 2 (outcome valences) x 3 (partners) ANOVA on average VS activation during the outcome phases of the Shared Reward task and Trust Game. B) Average VS activation (Z-statistics) during reward and loss sharing with each partner in the Shared Reward Task. VS activation was greatest during reward sharing with a friend but did not differ across partners during loss sharing. C) Average VS activation (Z-statistics) during reciprocation and defection from each partner in the Trust Game. VS activation was greatest when reciprocated by a friend but did not differ across partners during defection. Error bars reflect standard error; n = 123.

**Figure 4.**
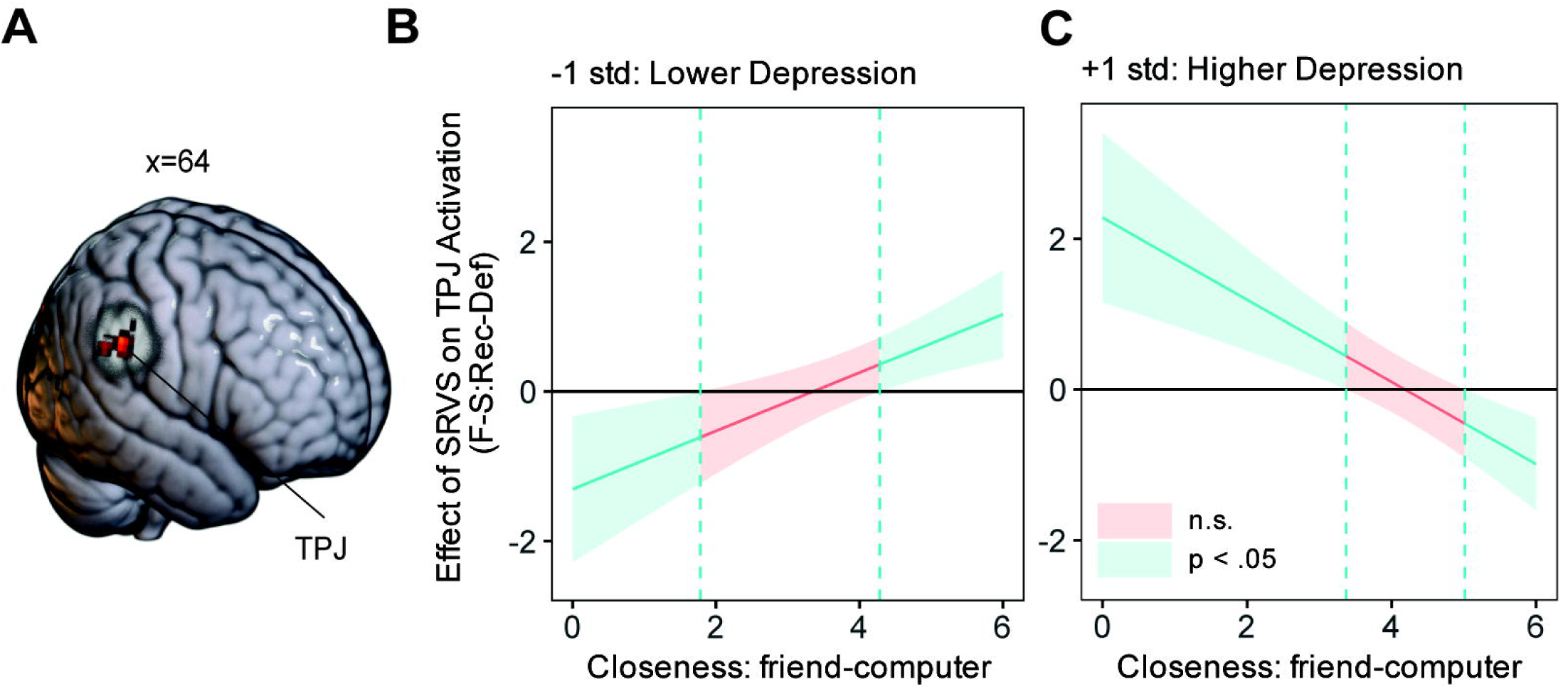
Depression reverses the association between striatal responsivity and TPJ activation during trust reciprocation. A) Whole-brain analysis of the friend–computer reciprocate > defect contrast revealed a significant three-way interaction in right TPJ, with striatal SRS as predictor and closeness and depressive symptoms as joint moderators. Clusters were thresholded at Z > 3.1 and corrected for multiple comparisons at p < .05. Thresholded and non-thresholded images are available on Neurovault (identifiers.org/neurovault.collection:23155) B) At low levels of depression (mean – 1 SD), striatal SRS positively predicted TPJ activation during reciprocation from close friends and negatively for distant friends. C) At high levels of depression (mean + 1 SD), this pattern reversed. Johnson–Neyman plots illustrate the conditional effect of striatal SRS on TPJ activation across closeness values; shaded cyan areas indicate significant effects (p < .05) and pink areas non-significant effects. Depression was modeled continuously in all analyses. Rec–Def = reciprocate > defect contrast.

**Figure 5.**
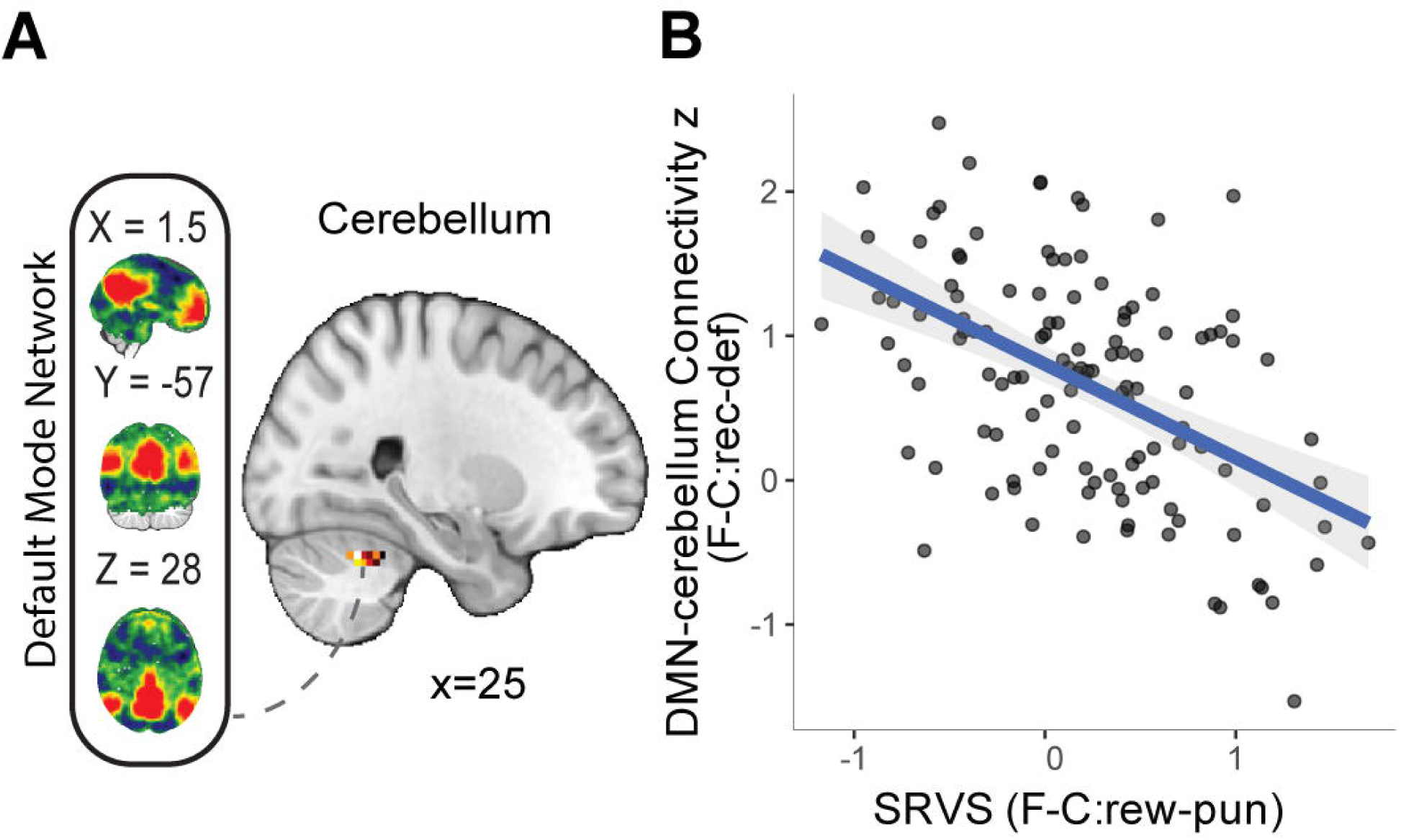
Striatal SRS negatively predicts DMN–cerebellum connectivity during trust reciprocation. A) Whole-brain network-based psychophysiological interaction analysis with DMN as the seed network revealed a cerebellar cluster in which connectivity with the DMN was negatively predicted by striatal SRS, based on the friend > computer reciprocate > defect contrast. Results were thresholded at Z > 3.1 and corrected for multiple comparisons at p < .05. Thresholded and non-thresholded images are available on Neurovault (identifiers.org/neurovault.collection:23155). B) Scatterplot illustrates the negative association between striatal SRS and DMN–cerebellum connectivity. F-C Rec–Def = friend > computer reciprocate > defect contrast; DMN = default mode network.

To identify additional regions where striatal SRS related to activity during trust reciprocity, we conducted whole-brain voxelwise analyses using the same models. Greater VS shared-reward sensitivity was associated with stronger activation to trust reciprocity in the superior parietal lobule (SPL, MNI = 33, −48, 41; cluster = 27 voxels, p = .013), and showed a closeness-dependent effect in retrosplenial cortex (RSC, MNI = −2, −53, 8; cluster = 24 voxels, p = .020), such that the association was positive for close friends and negative for distant friends. Whole-brain three-way interaction models revealed a broader pattern, including effects in TPJ, posterior cingulate/precuneus, dorsal lateral prefrontal cortex, and dorsal anterior cingulate cortex (Supplementary Table 2; Figure 3). In each of these regions, greater VS shared-reward sensitivity predicted weaker activation when participants high in depressive symptoms were reciprocated by a close friend, but stronger activation when reciprocation came from a more distant friend. The reverse pattern emerged among participants low in depressive symptoms. Supplementary analyses using the friend–stranger contrast revealed different activation patterns and interaction effects than those observed in the friend–computer analyses (see Supplement).

### 3.3 Does Striatal SRS Predict DMN Connectivity During Reciprocity?

We next tested whether individual differences in VS shared-reward sensitivity (Friend > Computer) predicted connectivity within the social valuation network during reciprocity. Specifically, we examined connectivity between a default mode network (DMN) seed and two preregistered targets—mPFC and VS—during reciprocate > defect outcomes from a friend relative to a computer partner. In separate models including either self-reported closeness or depressive symptoms as moderators, VS shared-reward sensitivity did not predict DMN–mPFC connectivity (closeness model: β = −0.120, SE = 0.225, t = −0.53, p = .59; depression model: β = –0.001, SE = 0.120, t = –0.01, p = .99) or DMN–VS connectivity (closeness model: β = 0.158, SE = 0.233, t = 0.68, p = .50; depression model: β = 0.057, SE = 0.124, t = 0.46, p = .65). Neither closeness nor depressive symptoms significantly moderated these associations. Exploratory models using the Friend > Stranger contrast yielded a similar pattern of null results (see Supplement).

Although ROI analyses showed no reliable associations between striatal SRS and DMN connectivity, whole-brain analyses revealed reduced DMN connectivity with the cerebellum in individuals with greater VS shared-reward sensitivity. Specifically, in models that separately examined closeness and depressive symptoms as moderators, greater striatal SRS was associated with reduced DMN connectivity in a cluster spanning cerebellar lobule V/VI (closeness model: MNI = 25, –45, –27; 33 voxels, p = .005; depression model: MNI = 25, –53, –27; 25 voxels, p = .019). Exploratory friend–stranger analyses further revealed a striatal SRS x closeness x depression three-way interaction in DMN–VS connectivity and a depression-related striatal SRS effect on DMN’s coupling with the dorsolateral prefrontal cortex (see Supplemental Figure 2). Together, these findings suggest that VS shared-reward sensitivity may relate to network-level responses during reciprocity, with aspects of this relationship varying as a function of depressive symptoms (see Supplement).

## 4. Discussion

It is widely understood that the VS tracks reward value and demonstrates sensitivity to social context. It has remained unclear, however, whether this neural correlate of social value is indicative of variations in behavior and brain responses across disparate social processing domains. Despite evidence that depression influences social reward processing, it remains unknown how altered VS signaling may shape engagement of specific brain systems and consequent behavior in social tasks. In this study, we used two tasks — the Shared Reward Task and the Trust Game — in which participants shared monetary outcomes with a close partner and had their trust either reciprocated or violated by that same partner. We considered whether an index of striatal sensitivity to social reward sharing (the striatal SRS) could predict behavior, regional activation, and/or interregional connectivity during the Trust Game.

Replicating earlier work, we found that participants reported greater closeness with friends, invested more in them, and displayed stronger striatal activity when sharing rewards with them. We also found that depression predicted relative investment in the friend, with greater depression coinciding with larger investment. We further observed that striatal SRS predicted closeness-dependent activation in SPL and RSC, regions implicated in social processing (Du et al., 2021; Shi et al., 2024). Further, jointly moderated by closeness and depression, striatal SRS predicted TPJ activation during friend’s (versus computer’s) reciprocity — a region known for mentalizing (Arioli et al., 2021), intent inference (Schurz et al., 2017), and impression updating in close relationships (Park et al., 2021).

We also observed an inverse relationship between striatal SRS and functional connectivity between the cerebellum and DMN during a friend’s (vs. computer’s) reciprocity. Though cerebellar function has historically been linked to motor control, recent work demonstrates that it also supports domain-general prediction and supervised learning (Manto et al., 2024; Van Overwalle et al., 2020), and that specific subregions are fundamental to social information processing (Chao et al., 2021; Pierce et al., 2023). This finding also accords with evidence that the cerebellum is anatomically (Schmahmann & Pandya, 1997) and functionally (Carta et al., 2019) coupled with cortico-striatal reward circuitry, particularly during social processing (Stoodley & Tsai, 2021). Together, these results suggest that stronger striatal SRS with a friend reduces information flow between DMN and cerebellar social learning circuitry during that friend’s reciprocation — perhaps because familiar partners don’t demand the same dynamic updating of social impressions. Confirmation will require follow-up studies examining how experimentally controlled variations in striatal signaling affect DMN-cerebellar connectivity and social valuation learning.

Together, these results suggest that the striatal social reward signal calibrates neural resources devoted to evaluating partner trustworthiness in accordance with perceived social closeness, aligning with an adaptive regulatory role of VS documented previously (e.g., Telzer, 2016), but that depression alters this association. In the case of higher depression, we observe elevated investment in closer friends, and an amplified striatal SRS that predicts TPJ hypo-activation in response to a close friend’s reciprocation but TPJ hyper-activation in response to a more distant friend’s reciprocation. This suggests that a depressed individual’s heightened valuation of shared reward with a close friend also dampens the neural response underlying evaluation of that friend’s trustworthiness. This complex pattern of amplified and dampened regional activity offers an explanation for the inconsistent observations of social blunting (Ait Oumeziane et al., 2019; Caouette & Guyer, 2016; Gradin et al., 2015) and sensitization (Gillard et al., 2021; Sankar et al., 2019; Steger & Kashdan, 2009; Yttredahl et al., 2018) under depression, showing that the two effects can coexist. It would be interesting to test whether this juxtaposition extends to striatal signaling in non-social value-based processes as well.

We acknowledge two key limitations. First, contrary to our predictions, neither continuous ratings of social closeness nor striatal SRS signal intensity directly predicted investment behavior. This absence may reflect learning-driven updates to perceived closeness and trust during the game, such that initial ratings are revised on the basis of actual outcomes (Daniel & Pollmann, 2014), obscuring any prior association with regional activation. Future experiments could assess closeness intermittently during gameplay or experimentally manipulate reciprocation rate to tease apart learning and initial valuation. Second, we assessed depressive symptomatology using the 7-up-7-down scale in a non-clinical sample, which may limit sensitivity to clinical-range depression and the ability to disentangle symptom domains such as anhedonia and depressive mood.

Despite these limitations, the current study provides the first evidence that VS reward sensitivity in one social context (reward sharing) predicts activation and connectivity in regions involved in social valuation in another social context (trust reciprocation). This cross-task prediction approach could be extended to other value-based social processes to test the generalizability and trait-like nature of striatal social sensitivity. Additionally, the current findings point to a potential neural mechanism that can produce both blunting and sensitization simultaneously, with the direction depending on the social processing demands of the task and the closeness of the partner. Together, these findings suggest that social dysfunction in depression may be nuanced, with sensitization in one social domain acting as a marker for blunting in another.

## Data and code availability

Analysis code and de-identified data related to this project can be found on GitHub (https://github.com/DVS-Lab/r01-soi) and OpenNeuro (https://openneuro.org/datasets/ds005123/).

## Acknowledgments

This work was supported in part by a grant from the National Institute on Aging (R01-AG067011 to DVS). This research includes calculations carried out on HPC resources supported by the US Army Research Laboratory under contract number ARL W911NF212007. We thank Cooper Sharp for assistance with data collection and data analysis, and Abraham Dachs, Arun Lakshmanan, Ashley Hawk, Derrick Dwamena, Hannah van Heerden, Ishika Kohli, Jamie-Nicole Luistro, Jenelle Scholl, Matthew McCormick, Melanie Kos, and Ryan Gephart for their contributions to data collection. A preprint version of this work is available on bioRxiv.

