## Supplement for "Striatal Social Reward Sensitivity Predicts Trust-Related Brain Responses Depending on Closeness and Depression"

### 1. Supplementary Methods

#### 1.1 Participants and Exclusion Criteria

We recruited 225 participants from the greater Philadelphia metro area using online and offline advertisements. The target sample size and exclusion criteria were preregistered (<https://osf.io/p654k>). Participants were screened to exclude major neurological illness and MRI contraindications, and all provided written informed consent in accordance with protocols approved by the Temple University IRB. Participants received $25 per hour for scanning sessions and $15 per hour for all other study procedures; they also were informed they would receive performance-based bonuses on one randomly selected trial per task to ensure incentive compatibility. The imaging and behavioral data used in the present analyses were collected as part of a larger project described in Smith et al. (2024) and are publicly available on OpenNeuro (<https://openneuro.org/datasets/ds005123/versions/1.1.3>).

Consistent with our preregistered analytic plan, we excluded 74 participants over age 55 to focus on a younger adult sample. Additional exclusions were applied for missing usable neuroimaging data (shared reward: n = 3; trust game: n = 4), missing individual difference measures (n = 7), and failed closeness manipulation (n = 8), defined as rating a computer or stranger as closer than one’s nominated friend. One participant was excluded for missing more than 25% of trust game trials, and we excluded an additional eight participants based on poor image quality using MRIQC metrics from echo-2 data (temporal SNR < Q1 − 1.5 x IQR or mean framewise displacement > Q3 + 1.5 x IQR). One final participant was excluded due to an experimental error (see deviation from preregistration). This yielded final samples of n = 123 for all analyses, whose demographic information is presented in Supplementary Table 1.

#### 1.2 Detailed Task Procedures

Participants performed two socio-economic tasks administered in PsychoPy (Peirce et al., 2019) in counterbalanced order, assessing neural responses to joint monetary outcomes in the Shared Reward Task and to partners’ reciprocity or defection in the Trust Game, both with social and nonsocial partners. Participants were asked to provide a real-life friend’s picture and were told their friend would participate in two tasks used in this study as one of their partners.

In each trial of the Shared Reward Task (Figure 1a), participants first saw which partner they would play with—friend, stranger, or computer—presented by name and photo. They then guessed whether a randomly generated hidden number would be greater or less than five. Feedback indicated the true number and whether the guess was correct: correct guesses earned both the participant and partner $10, incorrect guesses led to a $5 loss for both, and a draw occurred when the number was five. Trial outcomes were randomly generated to produce equal numbers of win and loss trials (each 4/9 of total) and fewer neutral trials (1/9).

In each trial in the Trust Game (Figure 1b), the participants again played with one of the same three types of partners: the same friend, stranger, or computer. The participants were endowed with $8 and could choose to invest less or more in the partner. The money invested in the partner was tripled, and the partner may choose to reciprocate by returning half of that tripled amount to the participant, or defect by keeping all of it. These either resulted in the participant gaining 1.5 of the amount they invested, or losing all of it. All partners were programmed to have a 50/50 chance of reciprocation. The Trust Game also contained two runs with 42 trials in total and 14 trials per partner in each run. For more details of the study procedures, see Smith et al. (2024).

#### 1.3 Behavioral Analyses

To replicate the “friend>stranger>computer” pattern of perceived closeness of and investment in partners (Fareri et al., 2012, 2015) we analyzed the IOS rating of and trial average investment in the three partners based on the Trust Game behavioral sample (n=123) using mixed-effects linear regression models with random intercepts. Additionally, to test whether striatal social reward sensitivity (SRS) cross-task predicts trust investment behavior, we regress trial average investment to the friend (vs other partners) on striatal SRS.

#### 1.4 Neuroimaging Data Preprocessing

All imaging data were first converted to BIDS format using HeuDiConv (Halchenko et al., 2024) prior to preprocessing. The results reported in this manuscript are based on data processed with fMRIPrep 24.1.1 (Esteban et al., 2019; Markiewicz et al., 2026), which builds on the Nipype framework (version 1.8.6, Esteban et al., 2025; Gorgolewski et al., 2011). Following developer guidelines, portions of the boilerplate text below are drawn directly from the fMRIPrep output. For clarity and relevance, we have removed extraneous details unrelated to our specific preprocessing steps.

**Anatomical data preprocessing.**

The T1w image was corrected for intensity non-uniformity (INU) with N4BiasFieldCorrection (Tustison et al., 2010), distributed with ANTs 2.5.3 (Avants et al., 2008), and used as T1w-reference throughout the workflow. The T1w-reference was then skull-stripped with a Nipype implementation of the antsBrainExtraction.sh workflow (from ANTs), using OASIS30ANTs as target template. Brain tissue segmentation of cerebrospinal fluid (CSF), white-matter (WM) and gray-matter (GM) was performed on the brain-extracted T1w using FAST (FSL 6.0.4, Zhang et al., 2001). Volume-based spatial normalization to two standard spaces (MNI152NLin6Asym, MNI152NLin2009cAsym) was performed through nonlinear registration with antsRegistration (ANTs 2.5.3), using brain-extracted versions of both T1w reference and the T1w template. The following templates were selected for spatial normalization and accessed with TemplateFlow (24.2.0,Ciric et al., 2022): FSL’s MNI ICBM 152 non-linear 6th Generation Asymmetric Average Brain Stereotaxic Registration Model (TemplateFlow ID: MNI152NLin6Asym, Evans et al., 2012), ICBM 152 Nonlinear Asymmetrical template version 2009c (TemplateFlow ID: MNI152NLin2009cAsym, Fonov et al., 2009).

**Functional data preprocessing.**

Then for all functional runs per participant, the following preprocessing was performed. First, a reference volume was generated from the shortest echo of the BOLD run, using a custom methodology of fMRIPrep, for use in head motion correction. Head-motion parameters with respect to the BOLD reference (transformation matrices, and six corresponding rotation and translation parameters) are estimated before any spatiotemporal filtering using mcflirt (FSL, Jenkinson et al., 2002). The estimated fieldmap was then aligned with rigid-registration to the target EPI (echo-planar imaging) reference run. The field coefficients were mapped onto the reference EPI using the transform. The BOLD reference was then co-registered to the T1w reference using mri_coreg (FreeSurfer) followed by flirt (FSL, Jenkinson & Smith, 2001) with the boundary- with six degrees of freedom. Several confounding time-series were calculated based on the preprocessed BOLD: framewise displacement (FD), DVARS and three region-wise global signals. FD was computed using two formulations following Power (absolute sum of relative motions, Power et al., 2014) and Jenkinson (relative root mean square displacement between affines, Jenkinson et al., 2002) FD and DVARS are calculated for each functional run, both using their implementations in Nipype (following the definitions by Power et al., 2014). The three global signals are extracted within the CSF, the WM, and the whole-brain masks.

Additionally, a set of physiological regressors were extracted to allow for component-based noise correction (CompCor, Behzadi et al., 2007). Principal components are estimated after high-pass filtering the preprocessed BOLD time-series (using a discrete cosine filter with 128s cut-off) for the two CompCor variants: temporal (tCompCor) and anatomical (aCompCor). tCompCor components are then calculated from the top 2% variable voxels within the brain mask. For aCompCor, three probabilistic masks (CSF, WM and combined CSF+WM) are generated in anatomical space. The implementation differs from that of Behzadi et al. in that instead of eroding the masks by 2 pixels on BOLD space, a mask of pixels that likely contain a volume fraction of GM is subtracted from the aCompCor masks. This mask is obtained by thresholding the corresponding partial volume map at 0.05, and it ensures components are not extracted from voxels containing a minimal fraction of GM. Finally, these masks are resampled into BOLD space and binarized by thresholding at 0.99 (as in the original implementation). Components are also calculated separately within the WM and CSF masks. For each CompCor decomposition, the k components with the largest singular values are retained, such that the retained components’ time series are sufficient to explain 50 percent of variance across the nuisance mask (CSF, WM, combined, or temporal). The remaining components are dropped from consideration. The head-motion estimates calculated in the correction step were also placed within the corresponding confounds file. All resamplings can be performed with a single interpolation step by composing all the pertinent transformations (i.e. head-motion transform matrices, susceptibility distortion correction when available, and co-registrations to anatomical and output spaces). Gridded (volumetric) resamplings were performed using nitransforms, configured with cubic B-spline interpolation. Many internal operations of fMRIPrep use Nilearn 0.10.4 (Abraham et al., 2014), mostly within the functional processing workflow. Many internal operations of *fMRIPrep* use *Nilearn* 0.6.2, mostly within the functional processing workflow. For more details of the pipeline, see the section corresponding to workflows in *fMRIPrep*'s documentation (<https://fmriprep.readthedocs.io/en/latest/workflows.html>).

Further, we applied spatial smoothing with a 5mm full-width at half-maximum (FWHM) Gaussian kernel respectively to all the functional data using FMRI Expert Analysis Tool (FEAT) Version 6.0. 2, part of FSL (FMRIB’s Software Library, [www.fmrib.ox.ac.uk/fsl](http://www.fmrib.ox.ac.uk/fsl)). Non-brain removal using BET (S. M. Smith, 2002) and grand mean intensity normalization of the entire 4D dataset by a single multiplicative factor were also applied.

#### 1.5 Echo-Time Dependent Denoising

TE-dependence analysis was performed on input data using the tedana workflow (DuPre et al., 2021). An initial mask was generated from the first echo using nilearn's compute_epi_mask function. An adaptive mask was then generated using the dropout method(s), in which each voxel's value reflects the number of echoes with 'good' data. An adaptive mask was then generated using the dropout method(s), in which each voxel's value reflects the number of echoes with 'good' data. A two-stage masking procedure was applied, in which a liberal mask (including voxels with good data in at least the first echo) was used for optimal combination, T2*/S0 estimation, and denoising, while a more conservative mask (restricted to voxels with good data in at least the first three echoes) was used for the component classification procedure. A monoexponential model was fit to the data at each voxel using nonlinear model fitting in order to estimate T2* and S0 maps, using T2*/S0 estimates from a log-linear fit as initial values. For each voxel, the value from the adaptive mask was used to determine which echoes would be used to estimate T2* and S0. In cases of model fit failure, T2*/S0 estimates from the log-linear fit were retained instead. Multi-echo data were then optimally combined using the T2* combination method (Posse et al., 1999). Principal component analysis based on the PCA component estimation with a Moving Average (stationary Gaussian) process (Li et al., 2007) was applied to the optimally combined data for dimensionality reduction. The following metrics were calculated: kappa, rho, countnoise, countsigFT2, countsigFS0, dice_FT2, dice_FS0, signal-noise_t, variance explained, normalized variance explained, d_table_score. Kappa (kappa) and Rho (rho) were calculated as measures of TE-dependence and TE-independence, respectively. A t-test was performed between the distributions of T2*-model F-statistics associated with clusters (i.e., signal) and non-cluster voxels (i.e., noise) to generate a t-statistic (metric signal-noise_z) and p-value (metric signal-noise_p) measuring relative association of the component to signal over noise. The number of significant voxels not from clusters was calculated for each component. Independent component analysis was then used to decompose the dimensionally reduced dataset. The following metrics were calculated: countnoise, countsigFS0, countsigFT2, d_table_score, dice_FS0, dice_FT2, kappa, normalized variance explained, rho, signal-noise_t, variance explained. Kappa (kappa) and Rho (rho) were calculated as measures of TE-dependence and TE-independence, respectively. A t-test was performed between the distributions of T2*-model F-statistics associated with clusters (i.e., signal) and non-cluster voxels (i.e., noise) to generate a t-statistic (metric signal-noise_z) and p-value (metric signal-noise_p) measuring relative association of the component to signal over noise. The number of significant voxels not from clusters was calculated for each component. Next, component selection was performed to identify BOLD (TE-dependent) and non-BOLD (TE-independent) components using a decision tree.

This workflow used numpy (Van Der Walt et al., 2011) scipy (Virtanen et al., 2020), pandas (McKinney, 2010; The pandas development team, 2024), scikit-learn (Pedregosa et al., 2011), nilearn (Nilearn contributors et al., 2026), bokeh (Bokeh Development Team, 2018), matplotlib (Hunter, 2007), and nibabel (Brett et al., 2019). This workflow also used the Dice similarity index (Dice, 1945; Sorenson, 1948).

#### 1.6 Creation of B0 Field maps

We created static B_0_ field maps from the phase information in our multi-echo functional MRI (ME-fMRI) echo-planar imaging (EPI) data using the Multi-Echo DIstortion Correction (MEDIC) algorithm (version 0.1.1; <https://github.com/vanandrew/warpkit>) (Van et al., 2023). This method estimates voxelwise B_0_ inhomogeneity by modeling the phase of the MRI signal as a linear function of echo time (Jezzard & Balaban, 1995).To remove channel-specific phase offsets introduced during multi-coil image reconstruction, we applied Multi-Channel Phase Combination using 3D Simultaneous estimation (MCPC-3D-S), which derives a zero-time phase offset based on the unwrapped phase difference between the first two echoes (Eckstein et al., 2018). Phase unwrapping was then performed using Rapid Open-source Minimum-spanning-tree phase unwrapping (ROMEO), which jointly unwraps all echoes under the constraint that phase accumulates linearly with echo time, thereby improving stability and accuracy (Dymerska et al., 2021). The resulting unwrapped phase data were fitted using a magnitude-weighted least-squares approach to compute off-resonance field maps in units of hertz, which were then transformed to anatomical space for use in susceptibility distortion correction.

#### 1.7 Neuroimaging Analyses

Neuroimaging analyses including activation and connectivity were performed on FSL version 6.0.7 (Jenkinson et al., 2012; S. M. Smith et al., 2004) using general linear models with local autocorrelation (Woolrich et al., 2001). For the shared reward run-level activation model, the twelve regressors include nine outcome event types with three outcome types (reward sharing, loss sharing, and neutral) intersecting with three partner identities (duration = 0.6 seconds), and three decision event types with different partners (duration = reaction time), with the outcome phase as our focus. We define striatal SRS as the ventral striatum ROI average activation from the contrasts of reward sharing > loss sharing outcome with a friend vs other partners in shared reward task, constrained by the portion of the Oxford-GSK-Imanova Striatal Connectivity Atlas (<https://fsl.fmrib.ox.ac.uk/fsl/fslwiki/Atlases/striatumconn>).

For the Trust Game run-level activation model, we used nine event-related regressors. First, to isolate the outcome phase, we included the six outcome regressors with two outcome natures (reciprocation and defection onsets) intersecting with three partner identities (duration = 1 second). To control variance related to expectation and decision, we also added three decision regressors with event onsets differentiated by three partners identities (duration = reaction time).

To operationalize the task-dependent connectivity between DMN and major social reward processing regions in Trust Game, we also built a network-based psychophysiological interaction (nPPI) analysis model on the individual. DMN as the seed network is defined by the DMN map among the ten network maps in Smith et al., (2009). We extracted the time series of spatial regression components of these ten networks’ activity during trust Game outcomes using a dual regression approach (Nickerson et al., 2017), and nine nPPI regressors were obtained from multiplying the DMN time series and the nine task regressors. The regressors in the model consist of the original nine trust game event regressors in activation models, nine nPPI regressors, and all ten network time series, resulting in 28 regressors in total. All task-related regressors were convolved with a double-gamma hemodynamic response function.

Nuisance regressors for the first-level analyses of both tasks include six motion parameters (translations and rotations), the first six components generated from aCompCor (anatomical component-based correction, see functional data preprocessing in the supplement) that explains the most variance, non-steady state volumes, and framewise displacement (FD) across time. To further control for non-BOLD noise, TEDANA-derived components were added as additional nuisance regressors. High-pass filtering (128s cut-off) was achieved using a set of discrete cosine basis functions.

After combining runs with a fixed-effects model, we conducted group-level analyses in FSL FLAME Stage 1 (Woolrich et al., 2004). Analyses began with pre-registered ROI models and were followed by whole-brain analyses using the same contrasts. Our primary focus was on reciprocated versus defected outcomes with a friend relative to other partners in the Trust Game. Anatomical ROIs were defined a priori: medial prefrontal cortex (mPFC) from a Neurosynth association map (keyword “mPFC,” z > 8), temporoparietal junction (TPJ) from the Mars connectivity atlas, and ventral striatum (VS) from the Oxford–GSK–Imanova Striatal Connectivity Atlas. In the connectivity analyses, we examined coupling between the default mode network (DMN) and key social-valuation regions, using mPFC and VS as target masks for nPPI analyses. Across models, we tested whether striatal SRS predicted trust behavior, activation, and connectivity using linear models that included striatal SRS, closeness, depressive symptoms, and their interaction terms. We also tested two-way and exploratory three-way interactions to assess potential joint moderation. All group-level models included age, gender, temporal signal-to-noise ratio (tsnr), and mean framewise displacement (fd_mean) as covariates. Whole-brain z-statistic maps were thresholded at z > 3.1 with cluster-extent correction at p < 0.05 (Worsley, 2001). Brain imaging results are displayed using MRIcroGL (Rorden, 2025)

### 2. Supplementary Results

#### 2.1 Closeness and Investment Patterns Across Partners

As a manipulation check, we examined whether participants rated their partners’ closeness and allocated investments consistent with perceived relational closeness. Although participants who rated their friend as less close than the other partners were excluded, we analyzed the trust game behavioral sample (n=123) using mixed-effects linear regression models with random intercepts. Participants reported feeling closer to their friend than to a stranger (β = 3.65, SE = 0.130, t(244) = 28.13, p < .0001, Tukey-corrected), and also slightly but significantly closer to a stranger than to a computer (β = 0.325, SE = 0.130, t(244) = 2.51, p = .03, Tukey-corrected). In terms of trust game behavior, participants invested significantly more in their friend than in either the stranger (β = 1.048, SE = 0.093, t(244) = 11.27, p < .0001, Tukey-corrected) or the computer (β = 1.206, SE = 0.093, t(244) = 12.96, p < .0001, Tukey-corrected). Average investments did not differ between the stranger and computer conditions (β = 0.157, SE = 0.093, t(244) = –1.70, p = .21, Tukey-corrected).

#### 2.2 Enhancement of Striatal SRS with Close Partner

We first sought to replicate prior work showing enhanced striatal responses during shared rewards and reciprocation with close partners (Fareri et al., 2012; 2015). We assessed whether VS activation varied as a function of social closeness during the outcome phases of both tasks. The 2 (reward/loss sharing outcomes) x 3 (partners) repeated-measures ANOVA on VS activation in the shared reward task revealed significant main effects of both partner (F(2,244)=3.63, p =.029, $\eta_{G}^{2}$ =.006; Mauchly’s test p > .05) and trial outcome (reward/loss sharing) (F(1,122)=161.58, p < .0001, $\eta_{G}^{2}$ =.188), as well as a significant interaction effect (F(2,244)=6.55, p =.002, $\eta_{G}^{2}$ =.011; Mauchly’s test p > .05). Striatal SRS was highest with friends, followed by strangers, then computers (Figure 3a in main text), with a significant difference between friend and computer conditions (β = 0.282, SE = 0.081, t(732) = 3.47, p = .002; Tukey-corrected) but not between friend and stranger (β = 0.164, SE = 0.081, t(732)= 2.01, p = .11; Tukey-corrected). For the Trust Game, the 2 (reciprocity/defect outcomes) x 3 (partners) repeated-measures ANOVA indicated significant main effects of both partner (F(2,244)=8.38, p =.0003, $\eta_{G}^{2}$ =.017; Mauchly’s test p > .05) and outcome (F(1,122)=92.45, p < .0001, $\eta_{G}^{2}$ =.109) on VS activation. Reciprocity from a friend again evoked stronger VS activation relative to both the stranger (β = 0.25, SE = 0.075, t(732)= 3.42, p = .002; Tukey-corrected) and computer (β = 0.191, SE = 0.075, t(732)= 2.54, p = .030; Tukey-corrected; Figure 3b in main text) conditions. The VS exhibited deactivation in response to negative outcomes in both tasks, and the responses could not be distinguished across partners.

#### 2.3 Striatal SRS and Trust Behavior: Friend-Stranger

To assess sensitivity to relational closeness within the social domain, we repeated the preregistered models using the Friend–Stranger contrast. VS SRS was defined as the reward > loss difference in ventral striatal activation when sharing outcomes with a friend relative to a stranger. In the model including self‑reported closeness, depressive symptoms, and their interaction, striatal SRS was not associated with closeness (β = –0.043, SE = 0.054, t = -0.802, p = .42), and there was no moderation by depression (β = –0.008, SE = 0.009, t = –1.06, p = .29). In a reduced model excluding depression terms, higher closeness was linked to lower striatal SRS (β = –0.084, SE = 0.041, t = –2.06, p = .04), opposite to the preregistered prediction.

We also examined whether VS shared‑reward sensitivity (Friend–Stranger) predicted trust behavior. striatal SRS did not relate to differences in average investment to the friend versus the stranger (β = –0.111, SE = 0.158, t = –0.71, p = .48), and the association was not significantly moderated by closeness (β = 0.136, SE = 0.099, t = 1.38, p = .17). Neither closeness (β = 0.004, SE = 0.071, t = 0.05, p=.96) nor depressive symptoms (β = 0.027, SE = 0.021, t = 1.28, p = .20) showed main effects on investment. These results mirror the Friend–Computer analysis: striatal SRS did not robustly predict partner‑selective trust behavior.

#### 2.4 Striatal SRS and Activation During Reciprocity: Friend-Stranger

We next tested whether striatal SRS to closeness (Friend–Stranger) predicted activation during reciprocate > defect outcomes from the friend partner. ROI models showed no association between striatal SRS to closeness and activation in mPFC (closeness model: β = 0.114, SE = 0.253, t = –0.45, p = .65; depression model: β = –0.068, SE = 0.151, t = 0.45, p = .65) or TPJ (closeness model: β = 0.062, SE = 0.201, t = 0.31, p = .76; depression model: β = –0.142, SE = 0.120, t = –1.18, p = .24), and no two‑way or three‑way interactions reached significance.

Whole‑brain voxelwise analyses identified closeness‑dependent associations between striatal SRS to closeness and activation in left middle temporal gyrus (MNI = –51, –42, –1; cluster = 28 voxels, p = .010; Supplemental Figure 1) and right cerebellum Crus I (MNI = 31, –64, –33; cluster = 29 voxels, p = .008). In both regions, higher striatal SRS to closeness was associated with greater activation when reciprocation came from a more distant friend, with the association reversing for close friends.

#### 2.5 Closeness and Depression Moderate the Link Between Striatal SRS and DMN Connectivity: Friend-Stranger

We examined whether striatal SRS predicted functional connectivity between the default mode network (DMN) and either the ventral striatum (VS) or medial prefrontal cortex (mPFC) during reciprocated (vs. defect) outcomes from the friend partner. In preregistered ROI models, striatal SRS was not significantly associated with DMN–mPFC connectivity (closeness model: β = –0.221, SE = 0.179, t = –1.23, p = .22; depression model: β = –0.134, SE = 0.105, t = –1.27, p = .21) or DMN–VS connectivity (closeness model: β = 0.182, SE = 0.199, t = 0.92, p = .36; depression model: β = 0.046, SE = 0.118, t = 0.39, p = .70). Closeness and depression did not moderate these effects in two-way models.

However, a significant three-way interaction emerged in DMN–VS connectivity (β = –0.018, SE = 0.009, t = –2.11, p = .037), suggesting that the association between striatal SRS and DMN connectivity varied as a function of both closeness and depressive symptoms. Among more depressed individuals, striatal SRS was trending toward a negatively associated with DMN–VS connectivity when reciprocation came from a closer friend; the pattern reversed for those reporting less depressive symptoms, who showed a trend-level positive association under the same conditions (Supplemental Figure 2).

We next explored whole-brain patterns of DMN connectivity using the same moderation models. A cluster in the dorsal lateral prefrontal cortex (dlPFC, MNI = – 26, 58, 14; 27 voxels, p = .014) showed a significant interaction with depressive symptoms: among more depressed individuals, greater striatal SRS was associated with reduced DMN–dlPFC connectivity, whereas the reverse pattern emerged for those with lower depressive symptoms (Supplemental Figure 3). A three-way interaction model of striatal SRS, closeness, and depression yielded a cluster in the primary motor cortex (MNI – 41, -15, 59; 34 voxels, p = .004).

#### 2.6 Striatal Specificity in Cross-Task Prediction of Social Valuation Regions

To confirm that striatal SRS carries region-specific predictive information — rather than reflecting a general neural signal — we repeated all cross-task ROI-based models using primary visual cortex (V1) as a control region, added as an additional interaction term to differentiate ROI-specific effects. The three-way interaction among striatal SRS, depression, and closeness in predicting posterior TPJ activation during a friend's (vs. computer's) reciprocation was specific to VS (β = −0.043, SE = 0.017, t = −2.57, p = .011) and was not observed for V1. Similarly, the interaction between striatal SRS and depression level in predicting DMN–mPFC connectivity was VS-specific, both for the friend–computer contrast (β = 0.055, SE = 0.024, t = 2.30, p = .023) and the friend–stranger contrast (β = 0.040, SE = 0.019, t = 2.08, p = .039). These results indicate that shared-reward striatal SRS carries region-specific information for predicting activation and connectivity in social valuation regions during trust reciprocation, even where the main effect of striatal SRS on DMN–mPFC connectivity did not reach significance.

### 3. Supplemental Figures and Figure Captions

**Supplemental Table 1**

Demographic table of the sample used in the study

| Variables | Share Reward and Trust Brain (n=123) |
| --- | --- |
| Age |  |
| Mean ± SD | 32.34 (8.97) |
| Min-Max | 21.12-54.66 |
| Median (IQR) | 30.78 (13.21) |
| Gender (%) |  |
| Female | 78 (63.42%) |
| Male | 40 (32.52%) |
| Non-binary | 5 (4.07%) |
| Race (%) |  |
| Asian | 18 (14.63%) |
| Black or African American | 18 (14.63%) |
| Two or more races | 8 (6.50%) |
| White | 75 (60.98%) |
| Prefer not to respond | 4 (3.25%) |
| Ethnicity (%) |  |
| Hispanic or Latino | 109 (88.62%) |
| Not Hispanic or Latino | 10 (8.13%) |
| Ethnicity not described (Other) | 4 (3.24%) |

**Supplemental Table 2.**

Clusters with activation showing a significant three-way interaction of striatal social responsivity, closeness and depression during friend (vs. computer) trust reciprocation

| Region |  | |  | X | MNI  Y | Z | Voxels | Z-statistics (max) | p-value |
| --- | --- | --- | --- | --- | --- | --- | --- | --- | --- |
| Occipital fusiform gyrus |  |  | | 14.3 | -77.4 | -12.6 | 1485 | 5.14 | 0 |
| Precuneus /superior parietal lobule |  |  | | -1.9 | -34.2 | 52.7 | 358 | 5.47 | 2.53e-19 |
| Lateral occipital cortex |  |  | | -28.9 | -85.5 | 23 | 222 | 4.68 | 8.65e-14 |
| Lateral occipital cortex |  |  | | -47.8 | -66.6 | 2.25 | 114 | 4.92 | 1.69e-08 |
| Lateral occipital cortex |  |  | | 35.9 | -82.8 | -0.72 | 93 | 4.19 | 2.98e-07 |
| Dorsal anterior cingulate |  |  | | -1.9 | 9 | 43.8 | 79 | 4.62 | 1.91e-06 |
| Dorsal premotor |  |  | | 27.8 | -7.2 | 46.8 | 79 | 4.71 | 1.91e-06 |
| Temporal parietal junction |  |  | | 62.9 | -31.5 | 37.9 | 56 | 4.49 | 6.29e-05 |
| Ventral occipito-temporal region |  |  | | 25.1 | -50.4 | -12.6 | 36 | 4.06 | 0.002 |
| Dorsal lateral prefrontal cortex |  |  | | 46.7 | 11.7 | 32 | 25 | 3.95 | 0.0175 |
| Superior parietal lobule |  |  | | -26.2 | -53.1 | 55.7 | 22 | 4.81 | 0.0332 |
| Dorsal premotor |  |  | | -26.2 | -4.5 | 52.7 | 21 | 4.1 | 0.0413 |

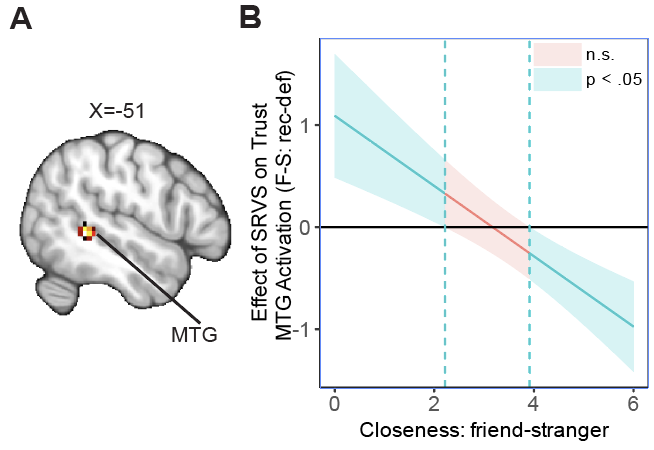

#### Supplemental Figure 1. Striatal social reward sensitivity (SRS) negatively predicts medial temporal gyrus (MTG) activation during close friends’ reciprocation.

**A**) Whole‑brain analyses identified a cluster in left MTG where activation during reciprocity (friend > stranger) was predicted by striatal activation, moderated by closeness. The result was thresholded and corrected for multiple comparisons using an initial cluster-forming threshold of Z > 3.1 followed by a whole-brain corrected cluster-extent threshold of p < 0.05. The thresholded and non-thresholded images can be accessed on Neurovault ( identifiers.org/neurovault.collection:23155). **B**) Johnson-Neyman plot demonstrates the conditional effect of striatal SRS on MTG activation during the reciprocation from friend vs stranger across levels of closeness. striatal SRS is negatively associated with MTG activation during close friend’s reciprocation but positively during distant friend’s. The Cyan sections suggest significant slopes (p<.05) at low and high levels of closeness and pink section non-significant slopes, with shaded area being the confidence interval. F-S: Rec-Def: Friend-stranger: reciprocation – defection.

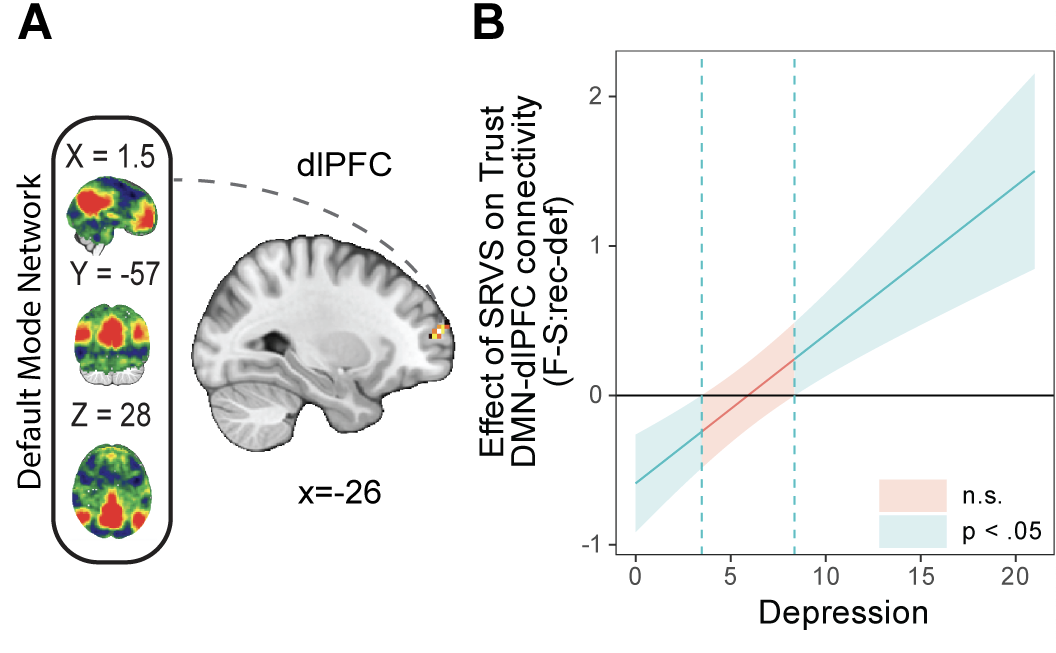

#### Supplemental Figure 2. Striatal social context sensitivity (striatal SRS) positively predicts DMN-dlPFC connectivity during friend’s reciprocation in participants with high depression symptomology.

**A**) We performed voxel-wise whole-brain psychophysiological interaction analyses using DMN as seed network on trust game’s outcome contrast of friend-stranger: reciprocate-defect, and located a left dlPFC cluster whose connectivity with DMN is predicted by striatal SRS and moderated by depression. The result was thresholded and corrected for multiple comparisons using an initial cluster-forming threshold of Z > 3.1. The thresholded and non-thresholded images can be accessed on Neurovault (identifiers.org/neurovault.collection:23155). **B**) Johnson-Neyman plot shows that among participants with higher depressive symptoms, greater striatal SRS predicted higher DMN–dlPFC connectivity during reciprocity, whereas this association is reversed in participants with lower symptoms. Cyan section suggests significant slopes (p<.05) at low and high levels of depression and pink section non-significant slopes, with shaded area being the confidence interval. F-S: Rec-Def: Friend-Stranger: Reciprocation-Defection, DMN: the default mode network, dlPFC: dorsal lateral prefrontal cortex.
